# N-terminal Domain Regulates Steroid Activation of Elephant Shark Glucocorticoid and Mineralocorticoid Receptors

**DOI:** 10.1101/822718

**Authors:** Yoshinao Katsu, Islam MD Shariful, Xiaozhi Lin, Wataru Takagi, Hiroshi Urushitani, Satomi Kohno, Susumu Hyodo, Michael E. Baker

## Abstract

Orthologs of human glucocorticoid receptor (GR) and human mineralocorticoid receptor (MR) first appear in cartilaginous fishes. Subsequently, the MR and GR diverged to respond to different steroids: the MR to aldosterone and the GR to cortisol and corticosterone. We report that cortisol, corticosterone and aldosterone activate full-length elephant shark GR, and progesterone, which activates elephant shark MR, does not activate elephant shark GR. However, progesterone inhibits steroid binding to elephant shark GR, but not to human GR. Together, this indicates partial functional divergence of elephant shark GR from the MR. Deletion of the N-terminal domain (NTD) from elephant shark GR (truncated GR) reduced the response to corticosteroids, while truncated and full-length elephant shark MR had similar responses to corticosteroids. Swapping of NTDs of elephant shark GR and MR yielded an elephant shark MR chimera with full-length GR-like increased activation by corticosteroids and progesterone compared to full-length elephant shark MR. Elephant shark MR NTD fused to GR DBD+LBD had similar activation as full-length MR, indicating that the MR NTD lacked GR-like NTD activity. We propose that NTD activation of human GR evolved early in GR divergence from the MR.

## Introduction

Glucocorticoids, such as cortisol and corticosterone, have diverse physiological activities in humans and other terrestrial vertebrates, including regulating glucose metabolism, cognition, immunity and the response to stress [1–5]. The physiological actions of glucocorticoids are mediated by the glucocorticoid receptor (GR), which belongs to the nuclear receptor family, a diverse group of transcription factors that also contains receptors for mineralocorticoids (MR), progestins (PR) androgens (AR) and estrogens (ER) [6–8].

The GR is closely related to the MR [9]; phylogenetic analysis indicates that the GR and MR evolved from a corticoid receptor (CR) that evolved in a jawless vertebrate in an ancestor of modern lamprey and hagfish [7,10–13]. Distinct orthologs of human GR and human MR first appear in cartilaginous fishes, the oldest group of extant jawed vertebrates (gnathostomes) that diverged from bony vertebrates about 450 million years ago [14]. Since the emergence of the GR and MR in cartilaginous fishes, these steroid receptors have diverged to respond to different corticosteroids. For example, human MR is activated by aldosterone, the physiological mineralocorticoid in humans, at a concentration at least ten-fold lower than by cortisol [15–18]. However, aldosterone has little activity for human GR, which is activated by cortisol and corticosterone (Figure 1) [6,12,16,19,20].

**Fig. 1.**
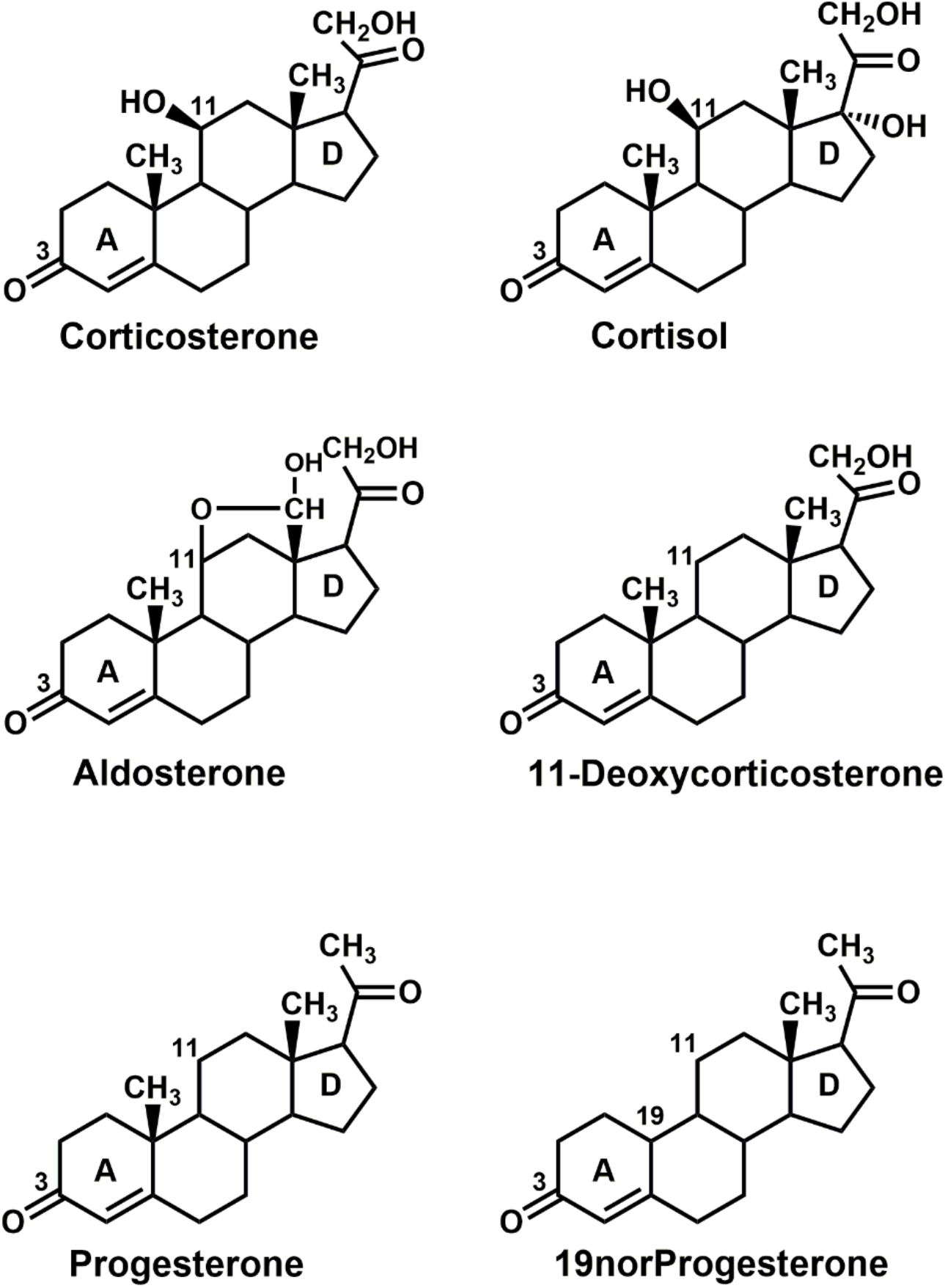
Structures of Corticosteroids and Progesterone. Cortisol and corticosterone are physiological glucocorticoids in terrestrial vertebrates and ray-finned fish [7,10,35]. Aldosterone, 11-deoxycorticosterone are physiological mineralocorticoids [10,12,36,37]. Progesterone is female reproductive steroid that also is important in male physiology [38,39].

The multi-domain structure of the GR is important in regulating steroid activation of the GR. Like other steroid receptors, the GR consists of an N-terminal domain (NTD) (domains A and B), a central DNA-binding domain (DBD) (domain C), a hinge domain (D) and a C-terminal ligand-binding domain (LBD) (domain E) [5,8,19,21,22] (Figure 2). The GR NTD is noteworthy for containing an activation function domain (AF1) that interacts with the DBD to provide a human GR NTD-DBD protein with constitutive activity [23–26]. AF1 also strongly stimulates transcriptional activation by glucocorticoids of the GR in humans [19,24,25,27], other terrestrial vertebrates [21,27–31] and ray-finned fish [21,32]. Interaction of the NTD with coactivators as well as post-transcriptional modification of the NTD also regulates transcriptional activation of the GR [5,22,30,33,34]. It is not known when strong activation by the NTD of vertebrate GRs evolved.

**Fig. 2.**
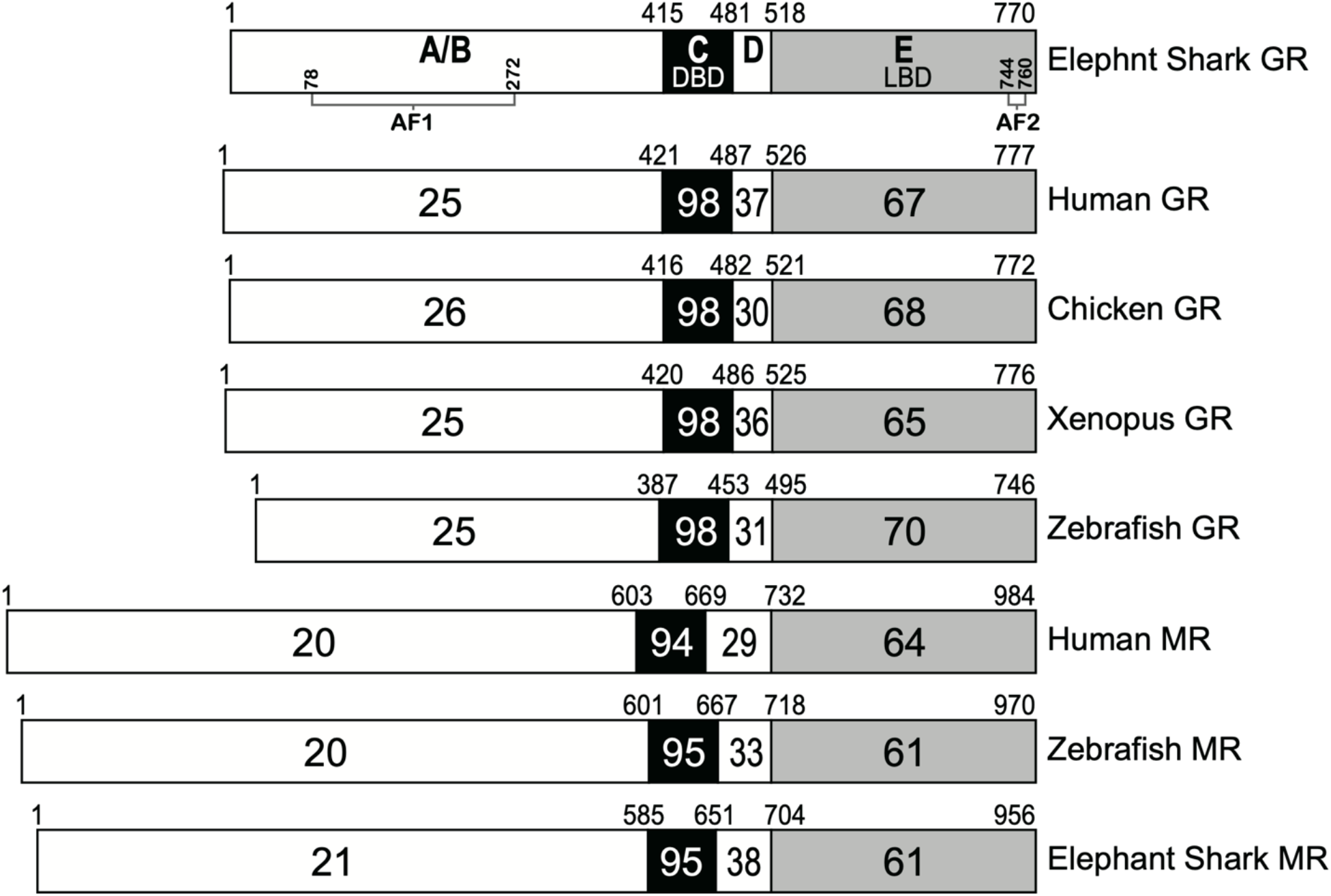
Comparison of domains in elephant shark GR with vertebrate GRs and MRs. GRs from elephant shark, zebrafish, *X. laevis*, chicken and humans and MRs from elephant shark, humans and zebrafish are compared. The functional NTD (A/B), DBD (C), hinge (D) and LBD (E) domains are schematically represented with the numbers of amino acid residues and the percentage of amino acid identity depicted. AF1 [5] and activation function 2 (AF2) [5] domains are shown in human GR.

To investigate corticosteroid signaling via the GR, in the light of the evolution [40], we decided to study activation of full-length and truncated elephant shark (*Callorhinchus milii*) GR by a panel of corticosteroids, which were used in a previous study of elephant shark MR [41].Although corticosteroid activation of another cartilaginous fish, skate GR, has been reported [42], this GR consisted of skate LBD fused to a GAL4-DBD. This GR lacked skate NTD and DBD, two domains that are important in AF1 activation of human GR [5,19,23–26]. The absence of data on steroid activation of a full-length cartilaginous fish GR prompted our investigation of the role of the NTD and DBD on transcriptional activation of a cartilaginous fish GR to determine if AF1 activity evolved in the NTD in a GR in a basal vertebrate. As the sequences of the NTDs of elephant shark and human GRs are poorly conserved, with 25% sequence identity (Figure 2), it was not evident that there would be strong AF1 activity in the NTD in elephant shark GR. We also investigated the evolution of GR specificity for glucocorticoids through a comparison of steroid activation of full-length elephant shark GR to elephant shark MR and human GR [16,19–21,23,27].

Here we report that full-length elephant shark GR is activated by cortisol and corticosterone, which activate human GR [16,19,20,27]. Unexpectedly, aldosterone and 11-deoxycorticosterone, two mineralocorticoids, also activated elephant shark GR. These two mineralocorticoids have little activity for human GR [15,16,27], which is selective for cortisol and corticosterone [16,20,27]. Elephant shark GR is not activated by either progesterone or 19norprogesterone, in contrast to elephant shark MR, which also is activated by cortisol and corticosterone [41]. However, we find that progesterone and 19norprogesterone inhibit corticosterone activation of elephant GR, while neither progestin inhibits cortisol activation of human GR.

We find that the NTD in elephant shark GR is a strong activator of steroid mediated transcriptional activation. Corticosteroid activation of truncated elephant shark GR, in which the NTD is deleted, was less than 10% of that of full-length GR, in contrast to truncated elephant shark MR, in which deletion of the NTD did not have a major effect on corticosteroid activation [41]. To investigate further the specificity of the each NTD for activation of the GR and MR, we studied chimeras of elephant shark GR and MR in which the GR NTD was fused to elephant shark MR DBD-LBD and the MR NTD was fused to GR DBD-LBD. In elephant shark GR NTD fused to MR DBD-LBD activation by cortisol and corticosterone increased by over 10-fold, compared to full length elephant shark MR indicating that this GR NTD contains AF1 activity. Interestingly, the chimera of GR NTD MR DBD-LBD had increased activation by progesterone and 19norprogestrone. In contrast, in MR NTD fused to GR DBD-LBD corticosteroid activation was reduced by over 90%. These data indicate allosteric regulation by the NTD evolved in elephant shark GR soon after the divergence of the GR from its common ancestor with the MR.

## Results

### Functional domains on elephant shark GR and other vertebrate GRs

In Figure 2, we compare the functional domains of elephant shark GR to corresponding domains in selected vertebrate GRs (human, chicken, Xenopus, zebrafish) and MRs (human, zebrafish and elephant shark). Elephant shark GR and human GR have 98% and 67% identity in DBD and LBD, respectively. Elephant shark GR and MR have 95% and 61% identity in DBD and LBD, respectively. This strong conservation of the DBD and LBD contrasts with the low sequence identity between the NTD in elephant shark GR and the NTD in other GRs and MRs. The NTD of elephant shark GR has only 21% sequence identity with elephant shark MR.

### Effect of corticosteroids and progesterone on transcriptional activation of full-length elephant shark GR and MR

We screened a panel of steroids (cortisol, corticosterone, 11-deoxycorticosterone, aldosterone, progesterone, 19norProgesterone) at 10^−8^ M for activation of full-length elephant shark GR (Figure 3A). For comparison, activation of full-length elephant shark MR by these steroids is shown in Figure 3B. Cortisol, corticosterone, 11-deoxycorticosterone, and aldosterone activated full-length elephant shark GR (Figure 3A) and MR (Figure 3B). However, fold-activation of full-length elephant shark GR by corticosteroids at 10^−8^ M was over 10-fold higher than for full-length elephant shark MR. For example, 10^−8^ M corticosterone activated full-length GR by 225-fold and full-length MR by 8-fold. Interestingly, neither progesterone nor 19norProgesterone activated elephant shark GR, while these steroids activated elephant shark MR, as previously reported [41].

**Fig. 3.**
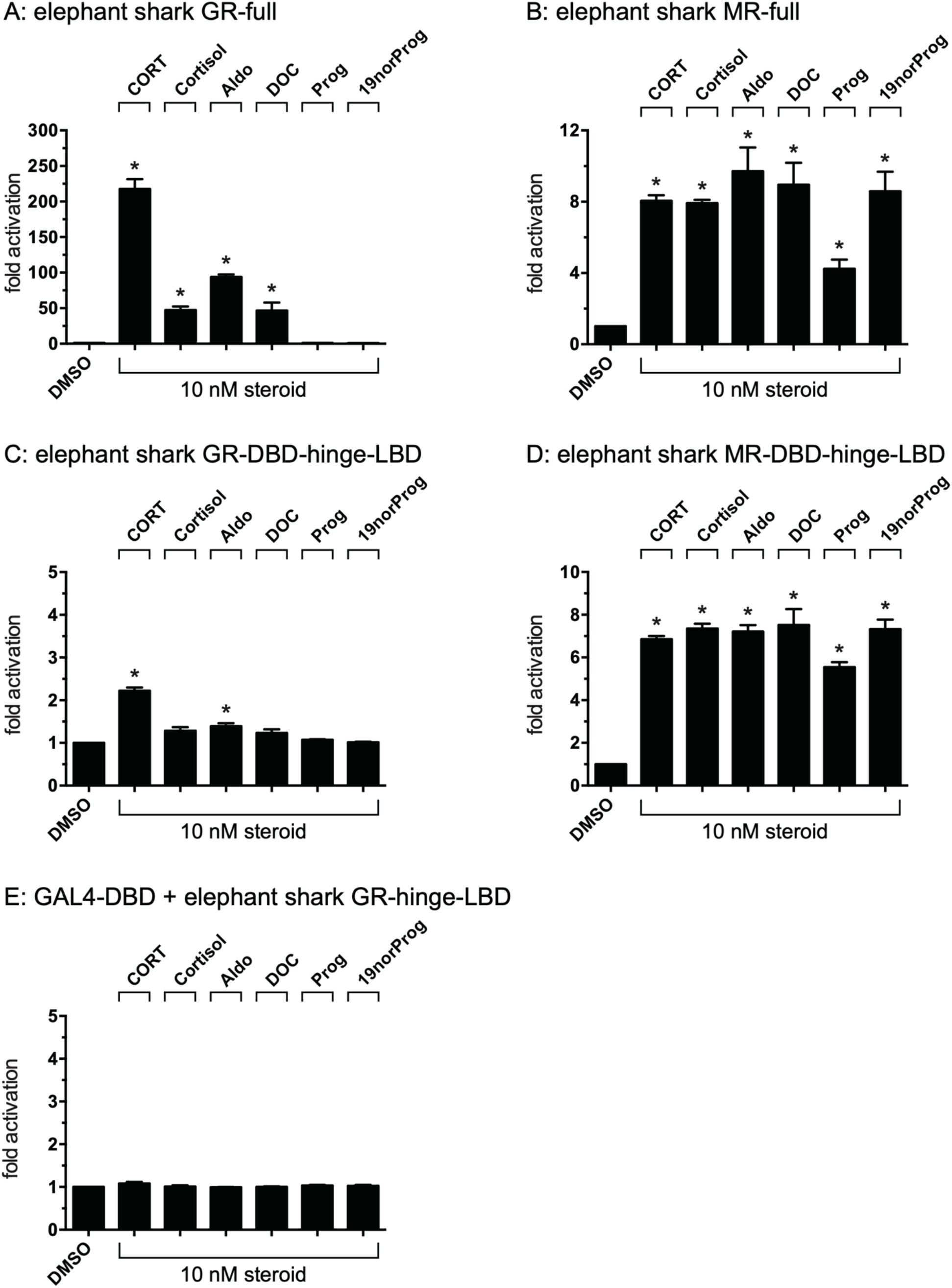
Ligand specificity of full-length and truncated elephant shark GR and MR. Plasmids for full-length elephant shark GR or MR or truncated elephant GR or MR, containing the DBD (C domain), D domain and LBD (E domain), were expressed in HEK293 cells with an MMTV-luciferase reporter. Transfected cells were treated with either 10 nM cortisol, corticosterone (CORT), aldosterone (Aldo), 11-deoxycorticosterone (DOC), progesterone (Prog), 19norProgesterone (19norProg) or vehicle alone (DMSO). Results are expressed as means ± SEM, n=3. Y-axis indicates fold-activation compared to the activity of control vector with vehicle (DMSO) alone as 1. A. Full-length elephant shark GR. B. Full-length elephant shark MR. C. Truncated elephant shark GR. D. Truncated elephant shark MR. E. GAL4-DBD+elephant shark GR-hinge-LBD.

### Effect of corticosteroids and progesterone on transcriptional activation of truncated elephant shark GR and MR

We investigated the role of the NTD in the response to corticosteroids and progestins at 10^−8^ M, of truncated elephant shark GR and MR, in which the NTD was deleted (Figure 3C, D). Figure 3C shows that activation of truncated elephant shark GR by 10^−8^ M corticosterone decreased by over 90%, compared to full-length elephant shark GR (Figure 3A), and there was no significant activation of truncated GR by 10^−8^ M of either cortisol or 11-deoxycorticosterone, both of which activated full-length GR. This indicates that NTD in elephant shark GR contains an AF1 domain. Progesterone did not activate truncated elephant shark GR. In contrast, at 10^−8^ M, corticosteroids and progesterone had similar levels of activation of truncated elephant shark MR (Figure 3D) and full-length MR (Figure 3B), indicating that there is a major difference between NTD activation of transcription of elephant shark GR and MR.

To investigate the effect of the elephant shark DBD on activation of elephant shark GR and MR, we replaced the DBD with GAL4 DBD, which has been used for analysis of transcription of the GR and MR [27,28,42,43]. As shown in Figure 3E, there was only a low level of activation by 10 nM corticosterone and aldosterone of GAL4 DBD GR hinge-LBD, and no activation by 11-deoxycorticosterone and cortisol. In contrast, GAL4 DBD MR LBD is activated by aldosterone, 11-deoxycorticosterone and other corticosteroids [41].

### Concentration-dependent activation by corticosteroids and progesterone of full-length and truncated elephant shark GR and MR

To gain a quantitative measure of NTD activation of elephant shark GR, we determined the concentration dependence of transcriptional activation by corticosteroids and progesterone of full-length and truncated elephant shark GR (Figure 4A, B) and for comparison of elephant shark MR (Figure 4C, D). This data was used to calculate a half maximal response (EC50) for steroid activation of elephant shark GR and MR (Tables 1 and 2). The graphs for cortisol, aldosterone and 11-deoxycorticosterone for elephant shark GR do not saturate. Thus, the EC50 values for these steroids shown in Table 1 are approximate and underestimate the true EC50. Corticosterone and cortisol, two physiological glucocorticoids in mammals, had EC50s of 7.9 nM and approximately 35 nM, respectively, for full-length elephant shark GR (Table 1). For comparison, the EC50s of corticosterone and cortisol are 0.61 nM and 1.6 nM, respectively, for full-length elephant shark MR (Table 2) [41]. However, aldosterone and 11-deoxycorticosterone, two physiological mineralocorticoids in mammals, had EC50s of approximately 24 nM and 28 nM, respectively, for elephant shark GR. Activation by these steroids of elephant shark GR indicates that it retains some properties of its MR paralog. For comparison, the EC50s of aldosterone and 11-deoxycorticosterone are 0.14 nM and 0.1 nM, respectively, for full-length elephant shark MR (Table 2) [41].

**Fig. 4.**
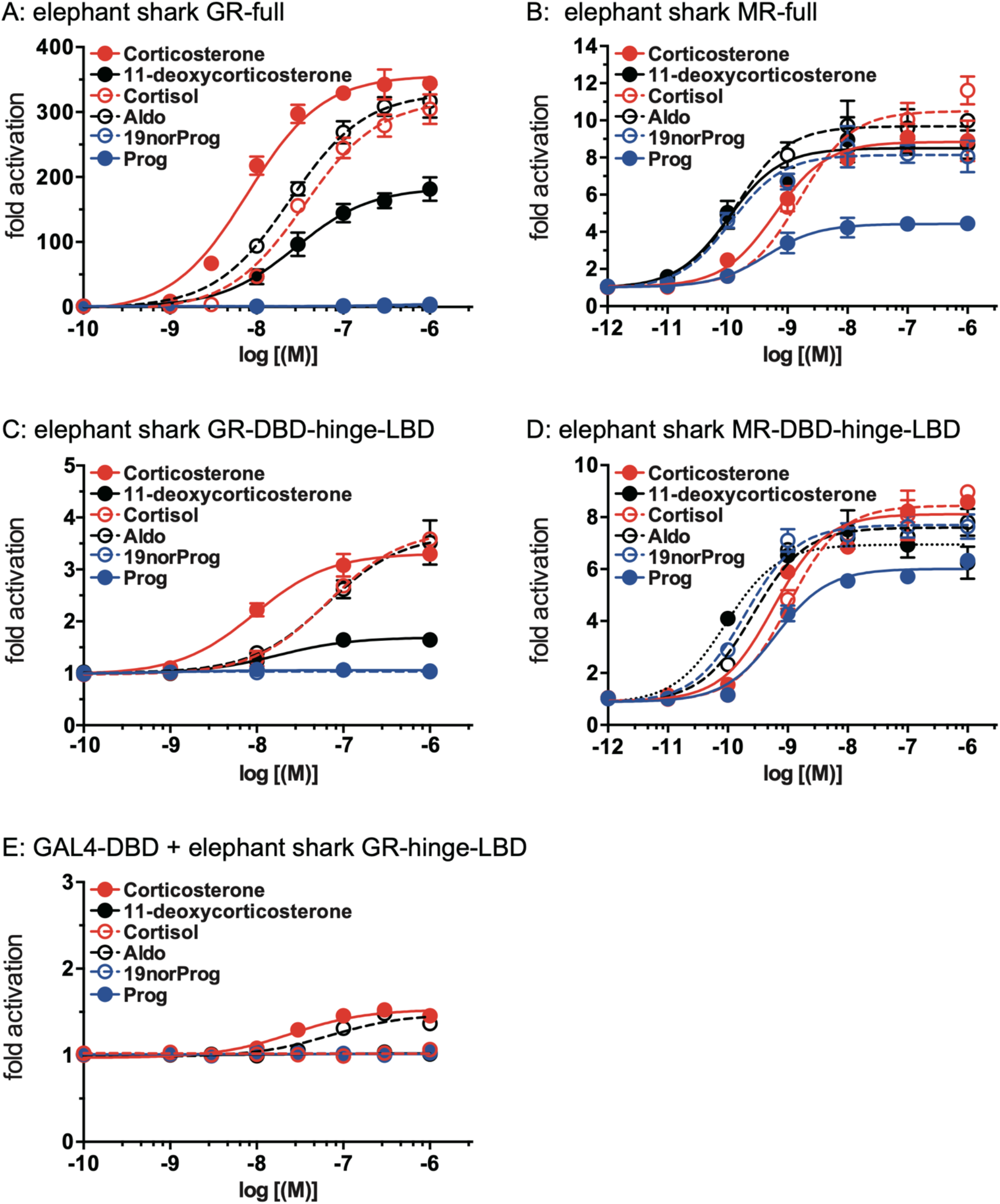
Concentration-dependent transcriptional activation by corticosteroids and progesterone of full length and truncated elephant shark GR and MR. Plasmids for full-length elephant shark GR or MR or truncated elephant GR or MR, were expressed in HEK293 cells with an MMTV-luciferase reporter. Cells were treated with increasing concentrations of either cortisol, corticosterone, aldosterone, 11-deoxycorticosterone, progesterone, 19norProgesterone or vehicle alone (DMSO). Results are expressed as means ± SEM, n=3. Y-axis indicates fold-activation compared to the activity of control vector with vehicle (DMSO) alone as 1. A. Full-length elephant shark GR. B. Full-length elephant shark MR. C. Truncated elephant shark GR (Domains CDE). D. Truncated elephant shark MR (Domains CDE). E. GAL4-DBD+elephant shark GR-hinge-LBD.

**Table 1.**
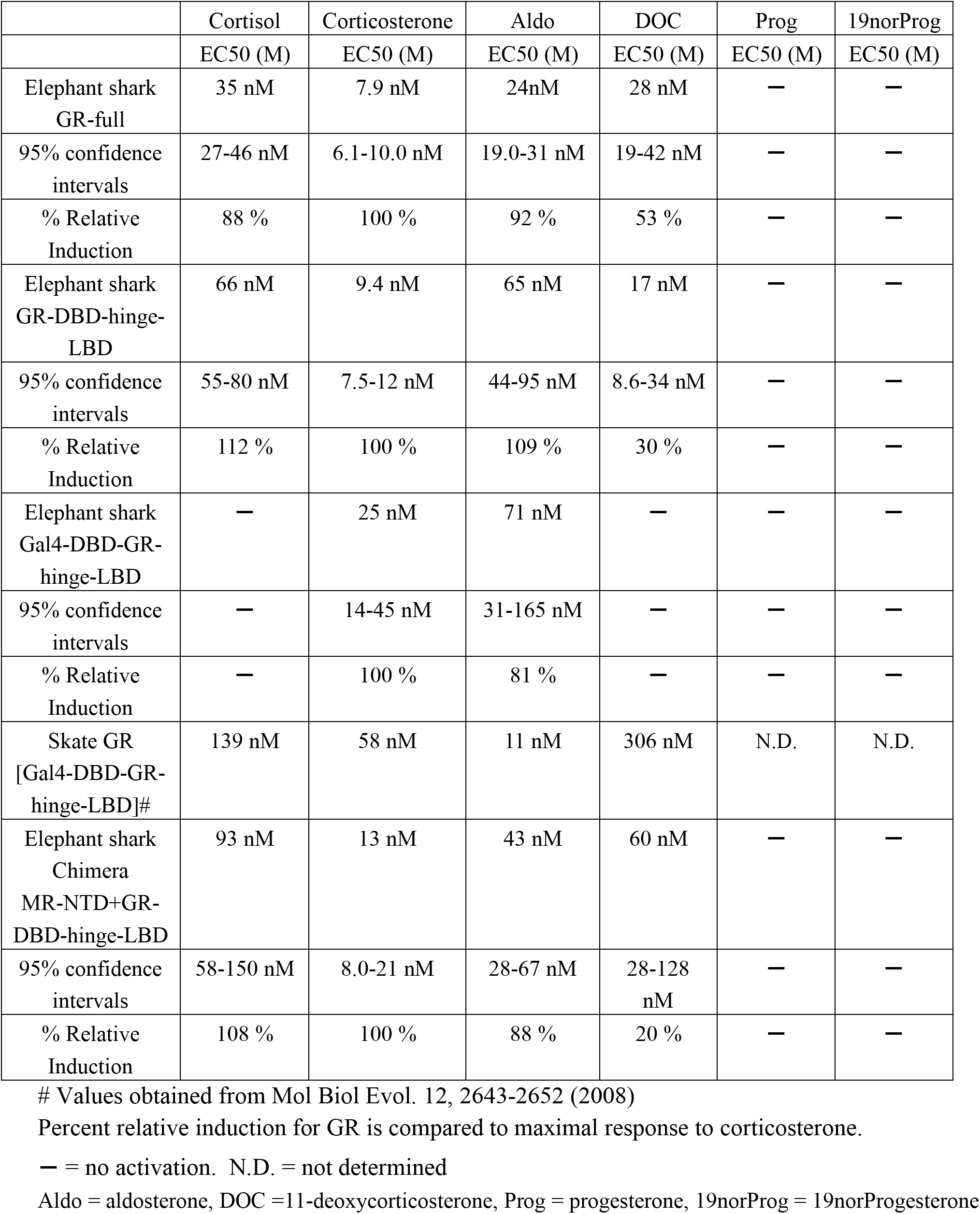
EC50 values for steroid activation of full-length and truncated elephant shark GR

|  | Cortisol | Corticosterone | Aldo | DOC | Prog | 19norProg |
| --- | --- | --- | --- | --- | --- | --- |
|  | EC50 (M) | EC50 (M) | EC50 (M) | EC50 (M) | EC50 (M) | EC50 (M) |
| Elephant shark GR-full | 35 nM | 7.9 nM | 24nM | 28 nM | — | — |
| 95% confidence intervals | 27-46 nM | 6.1-10.0 nM | 19.0-31 nM | 19-42 nM | — | — |
| % Relative Induction | 88 % | 100 % | 92 % | 53 % | — | — |
| Elephant shark GR-DBD-hinge-LBD | 66 nM | 9.4 nM | 65 nM | 17 nM | — | — |
| 95% confidence intervals | 55-80 nM | 7.5-12 nM | 44-95 nM | 8.6-34 nM | — | — |
| % Relative Induction | 112 % | 100 % | 109 % | 30 % | — | — |
| Elephant shark Gal4-DBD-GR-hinge-LBD | — | 25 nM | 71 nM | — | — | — |
| 95% confidence intervals | — | 14-45 nM | 31-165 nM | — | — | — |
| % Relative Induction | — | 100 % | 81 % | — | — | — |
| Skate GR [Gal4-DBD-GR-hinge-LBD]# | 139 nM | 58 nM | 11 nM | 306 nM | N.D. | N.D. |
| Elephant shark Chimera MR-NTD+GR-DBD-hinge-LBD | 93 nM | 13 nM | 43 nM | 60 nM | — | — |
| 95% confidence intervals | 58-150 nM | 8.0-21 nM | 28-67 nM | 28-128 nM | — | — |
| % Relative Induction | 108 % | 100 % | 88 % | 20 % | — | — |

**Table 2.**
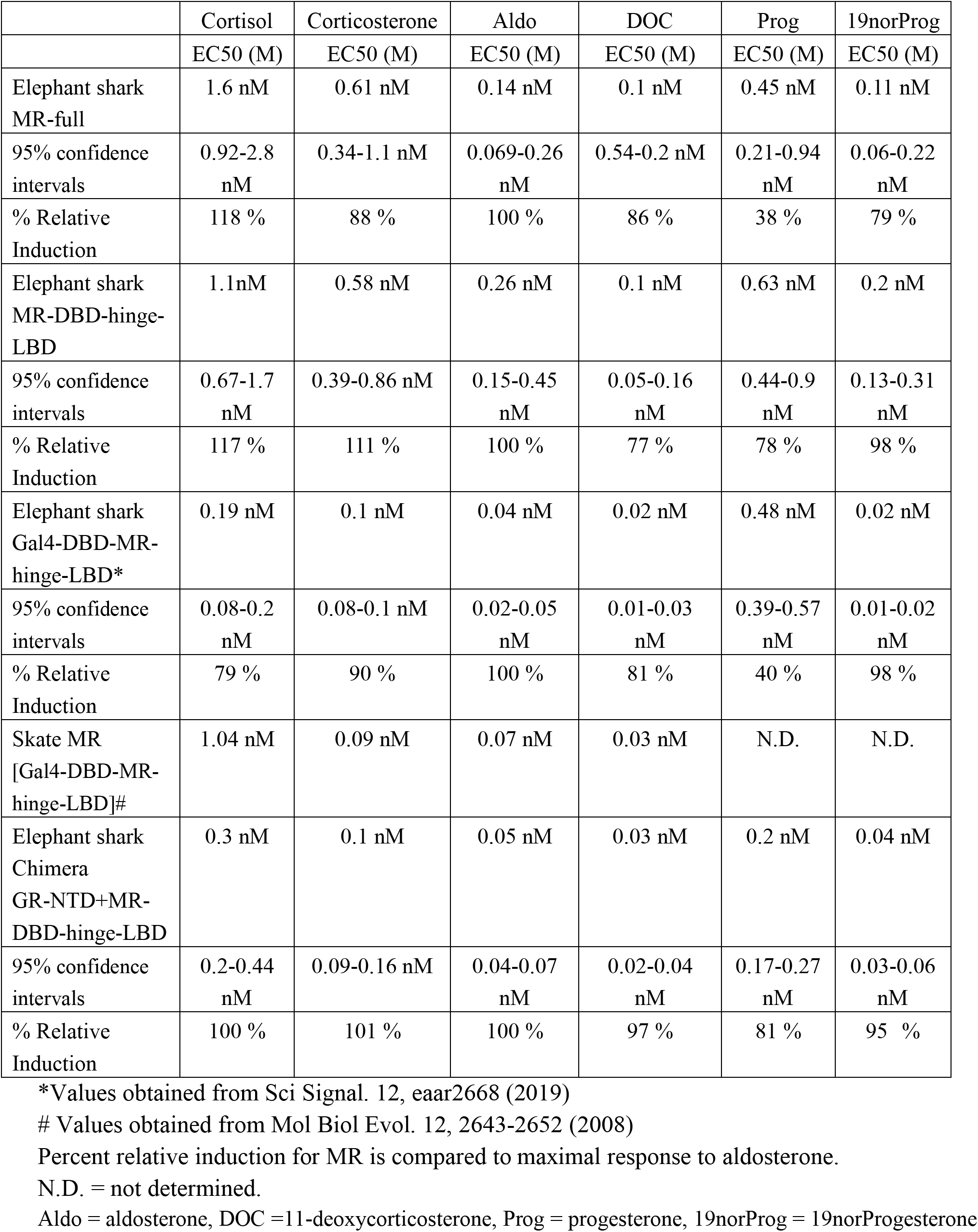
EC50 values for steroid activation of full-length and truncated elephant shark MR

|  | Cortisol | Corticosterone | Aldo | DOC | Prog | 19norProg |
| --- | --- | --- | --- | --- | --- | --- |
|  | EC50 (M) | EC50 (M) | EC50 (M) | EC50 (M) | EC50 (M) | EC50 (M) |
| Elephant shark MR-full | 1.6 nM | 0.61 nM | 0.14 nM | 0.1 nM | 0.45 nM | 0.11 nM |
| 95% confidence intervals | 0.92-2.8 nM | 0.34-1.1 nM | 0.069-0.26 nM | 0.54-0.2 nM | 0.21-0.94 nM | 0.06-0.22 nM |
| % Relative Induction | 118 % | 88 % | 100 % | 86 % | 38 % | 79 % |
| Elephant shark MR-DBD-hinge-LBD | 1.1 nM | 0.58 nM | 0.26 nM | 0.1 nM | 0.63 nM | 0.2 nM |
| 95% confidence intervals | 0.67-1.7 nM | 0.39-0.86 nM | 0.15-0.45 nM | 0.05-0.16 nM | 0.44-0.9 nM | 0.13-0.31 nM |
| % Relative Induction | 117 % | 111 % | 100 % | 77 % | 78 % | 98 % |
| Elephant shark Gal4-DBD-MR-hinge-LBD* | 0.19 nM | 0.1 nM | 0.04 nM | 0.02 nM | 0.48 nM | 0.02 nM |
| 95% confidence intervals | 0.08-0.2 nM | 0.08-0.1 nM | 0.02-0.05 nM | 0.01-0.03 nM | 0.39-0.57 nM | 0.01-0.02 nM |
| % Relative Induction | 79 % | 90 % | 100 % | 81 % | 40 % | 98 % |
| Skate MR [Gal4-DBD-MR-hinge-LBD]# | 1.04 nM | 0.09 nM | 0.07 nM | 0.03 nM | N.D. | N.D. |
| Elephant shark Chimera GR-NTD+MR-DBD-hinge-LBD | 0.3 nM | 0.1 nM | 0.05 nM | 0.03 nM | 0.2 nM | 0.04 nM |
| 95% confidence intervals | 0.2-0.44 nM | 0.09-0.16 nM | 0.04-0.07 nM | 0.02-0.04 nM | 0.17-0.27 nM | 0.03-0.06 nM |
| % Relative Induction | 100 % | 101 % | 100 % | 97 % | 81 % | 95 % |

\*Values obtained from Sci Signal. 12, eaar2668 (2019)

The EC50 of corticosterone for truncated elephant shark GR was 9.4 nM. The EC50s of cortisol and aldosterone for truncated elephant shark GR were approximately 66 nM and 65 nM respectively. In contrast, the EC50s of corticosterone and aldosterone for GAL4-DBD+GR-hinge-LBD were approximately 25 nM and 71 nM, respectively, and cortisol and 11-deoxycorticosterone did not activate GAL4-DBD+GR-hinge-LBD. The higher EC50 for corticosterone and the loss of activation by cortisol and 11-deoxycorticosterone, when elephant shark DBD is replaced with GAL4-DBD indicates that the DBD is important in transcriptional activation of elephant shark GR.

All four corticosteroids, progesterone and 19norProgesterone had EC50s for truncated MR (DBD+LBD) that were either similar to or lower than that of full-length MR (Table 2). EC50s for Gal4-DBD+MR-hinge-LBD were either similar to or lower than that of full-length MR.

The dependence of the level of cortisol and 11-deoxycorticosterone activation on the NTD in elephant shark GR (Figures 4A, C) is similar to that of human GR, in which deletion of the NTD reduces cortisol and 11-deoxycorticosterone fold-activation by over 90% [19,27]. The diminished activation of GAL4-DBD-elephant shark GR-hinge-LBD (Figure 4E) indicates that the DBD also is important in fold-activation of elephant shark GR. These results indicate that allosteric signaling between the NTD and DBD-LBD in elephant shark GR is critical for its response to corticosteroids. This contrasts to elephant shark MR, in which truncated MR and full-length elephant shark MR have similar levels of activation by corticosteroids and progesterone (Figures 4B, D). Together this indicates that an activation function in the GR NTD and higher EC50s for some corticosteroids that also activated elephant shark MR evolved early in the divergence of the GR from its MR kin.

### Progesterone binds to, but does not activate, full-length elephant shark GR

Neither progesterone nor 19norProgesterone activated transcription by elephant shark GR (Figure 3A), although these steroids are transcriptional activators of elephant shark MR (Figure 3B) [41]. The lack of progesterone activation of elephant shark GR could be due to decreased affinity of progesterone for elephant shark GR or to loss of activation of elephant shark GR after binding progesterone. We find that progesterone and 19norProgesterone inhibit activation of elephant shark GR by 10 nM corticosterone (Figure 5A) indicating that elephant shark GR recognizes progesterone. A parallel study showed that neither progesterone nor 19norProgesterone inhibited activation of human GR by 10 nM cortisol (Figure 5B). This indicates that the loss of activation by progesterone preceded the loss of progesterone binding to the GR, which occurred later in vertebrate evolution.

**Fig. 5.**
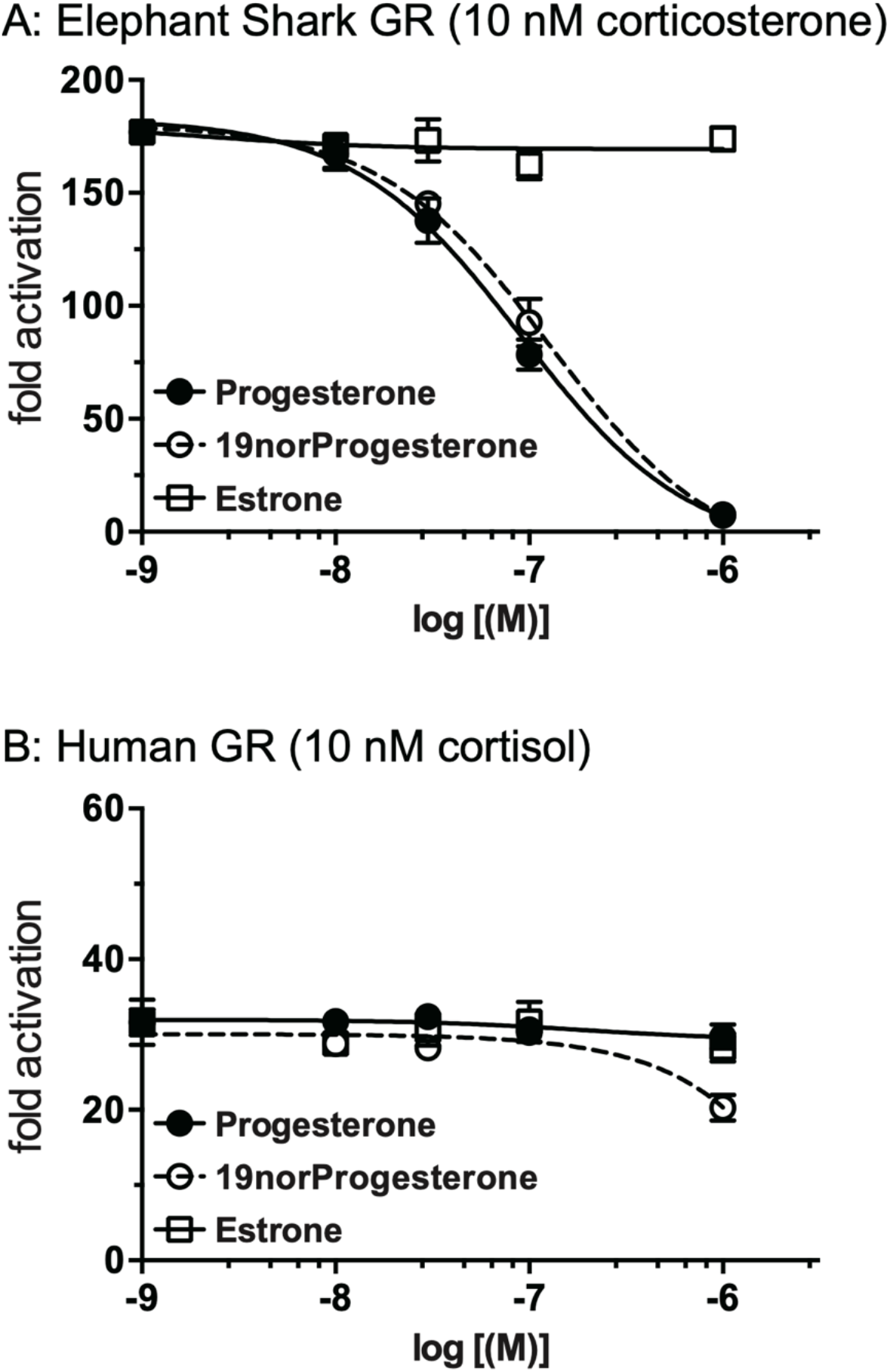
Progesterone inhibits corticosterone activation of full-length elephant shark GR. Plasmids encoding full length elephant GR or human GR were expressed in HEK293 cells. Cells with elephant shark GR and human GR were treated with 10 nM corticosterone and cortisol respectively and increasing concentrations of either progesterone, 19norProgesterone or estrone. Y-axis indicates fold-activation compared to the activity of control vector with vehicle (DMSO) alone as 1. A. Elephant shark GR. B. Human GR.

### Corticosteroid activation of chimeras in which the NTD is swapped between elephant shark GR and MR

To investigate further the origins of NTD activation of transcription by the GR, we studied corticosteroid activation of chimeras of elephant shark GR and MR, in which the GR NTD was fused to MR DBD-hinge-LBD and the MR NTD was fused to GR DBD-hinge-LBD (Figure 6). We find that despite low sequence similarity between the NTDs in elephant shark GR and MR, their NTDs had dominant effects on transcription of the DBD-hinge-LBD in each chimera (Figure 6, Tables 1 and 2). Thus, in the GR NTD-MR DBD-hinge-LBD chimera (GR NTD fused to MR DBD-hinge-LBD), cortisol and corticosterone increased activation by over 30-fold, compared to full length elephant shark MR (Figures 3B, 4B and 6B). Moreover, the GR NTD-MR DBD-hinge-LBD chimera also had a higher level of activation in the presence of progesterone and 19norProgestrone. In addition, the EC50s for corticosteroids and progestins were lower in the GR NTD-MR DBD-hinge-LBD chimera indicating that the NTD affects the affinity of steroids for the chimera, as well as the level of transcriptional activation (Table 2).

**Fig. 6.**
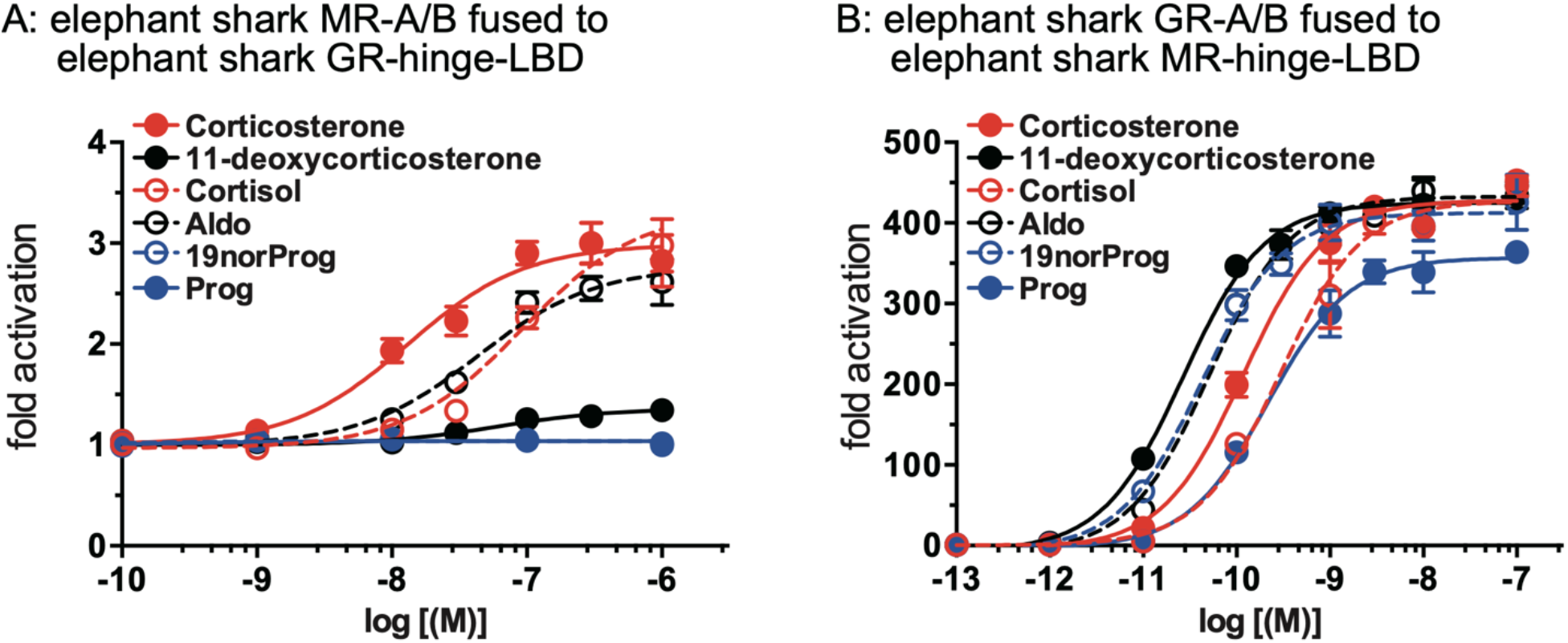
Concentration-dependent transcriptional activation by corticosteroids of chimeras of GR and MR. Plasmids for chimeric GR or MR were expressed in HEK293 cells with an MMTV-luciferase reporter. Cells were treated with increasing concentrations of either cortisol, corticosterone, aldosterone, 11-deoxycorticosterone, progesterone, 19norProgesterone or vehicle alone (DMSO). Results are expressed as means ± SEM, n=3. Y-axis indicates fold-activation compared to the activity of control vector with vehicle (DMSO) alone as 1. A. Chimera of MR-A/B fused to GR-DBD-hinge-LBD. B. Chimera of GR-A/B fused to MR-DBD-hinge-LBD.

In contrast, the MR NTD reduced activation by corticosterone and other corticosteroids of the MR NTD-GR DBD-hinge-LBD chimera by over 90% compared to full-length GR (Figures 3A, 4A and 6A, Table 1). These data indicate that increased activation by the NTD in elephant shark GR evolved soon after it diverged from the MR.

### RU486 is a glucocorticoid antagonist of full-length elephant shark GR

To gain another measure of the similarities and differences between elephant shark GR and human GR, we investigated the effect of RU486 on transcriptional activation of elephant shark GR. RU486, originally developed as a progesterone antagonist [44,45] also is used as an antagonist for glucocorticoid activation of human GR [44], although at high concentrations (>100nM), RU486 is a weak GR agonist. We found that RU486 inhibited activation by 2 nM dexamethasone of elephant shark and human GR (Figure 7A, B). At 0.9 nM, RU486 inhibited dexamethasone activation of human GR by 50% (Figure 7A). At 2.2 nM, RU486 inhibited dexamethasone activation of elephant shark GR by 50% (Figure 7B).

**Fig. 7.**
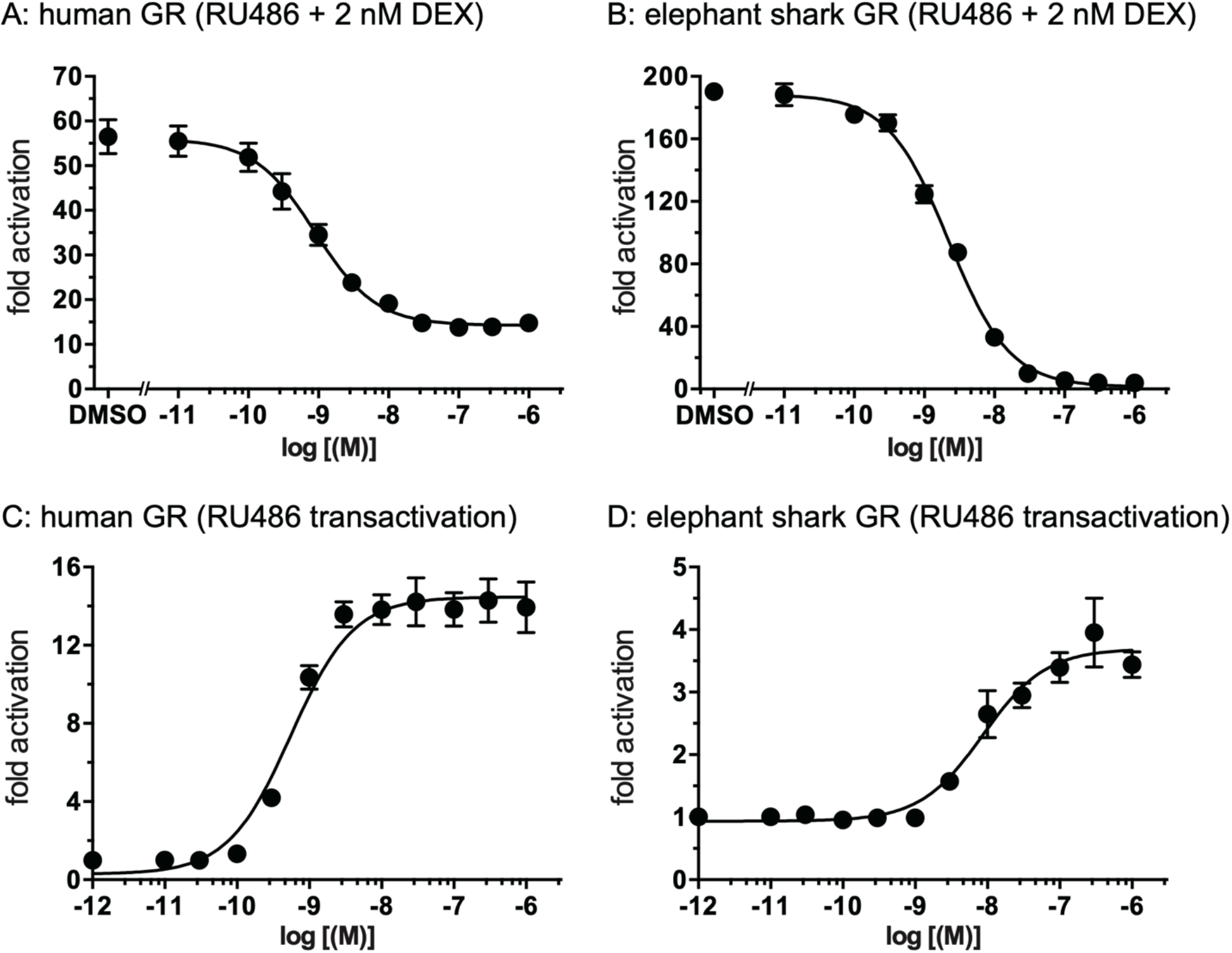
RU486 inhibition of dexamethasone activation of elephant shark GR and human GR. Plasmids encoding full length elephant GR or human GR were expressed in HEK293 cells. Cells with elephant shark GR or human GR were treated with 2 nM dexamethasone + increasing concentrations of RU486 (panels A, B) or increasing concentrations of RU486 (panels C, D). Results are expressed as means ± SEM, n=3. Y-axis indicates fold-activation compared to the activity of control vector with vehicle (DMSO) alone as 1.

Concentration dependent activation by RU486 of elephant shark GR and human GR, shown in Figure 7C and D, was used to calculate EC50 values for RU486. The EC50 of RU486 for elephant shark GR is 8.7 nM and for human GR is 0.55 nM. Maximal transcriptional activation by RU486 of elephant shark GR (Figure 7D) is considerably lower than for activation of human GR (Figure 7C). Nevertheless, although RU486 is not as active towards elephant shark GR as towards human GR, the responses of elephant shark GR to RU486 mimic those of human GR.

## Discussion

Sequence analysis revealed that the MR and GR are kin [9,13]. A distinct GR and MR first appear in cartilaginous fish, which occupy a key position at an ancestral node from which ray-finned fish and terrestrial vertebrates diverged [14,46,47]. Genomic analyses reveal that elephant shark genes are evolving slowly [14]. Thus, analysis of steroid activation of elephant shark GR can provide insights into the evolution of steroid activation of the ancestral GR early in its divergence from the MR. As a consequence, we studied activation of elephant shark GR by physiological glucocorticoids (cortisol, corticosterone) and mineralocorticoids (aldosterone, 11-deoxycorticosterone), as well as progesterone, which we compared to similar experiments with elephant shark MR [41]. Because the NTD has a strong AF1 for human GR [19,23–26], unlike human MR [19,41], we also studied elephant shark GR, in which the NTD was deleted. After finding that corticosterone activation of truncated elephant shark GR declined by over 90% from the activity of full-length elephant shark GR, we investigated the evolution of the NTD in chimeras of elephant shark GR and MR, in which their NTDs were swapped. Despite low sequence identity between the GR and MR NTDs, AF1 in elephant shark GR NTD could transfer its activity to elephant shark MR, when it was substituted for the MR NTD. This indicates that a strong AF1 appeared early in the evolution of the GR in a basal jawed vertebrate (Figure 6, Tables 1 and 2).

Overall, we find that the response to corticosteroids and RU486 [44,48] by elephant shark GR has some similarities and some differences to human GR [15,19,21,27], as well as to elephant shark MR [41]. In comparison to corticosteroid activation of elephant shark MR, corticosteroids have higher EC50s for elephant shark GR, indicating reduced sensitivity to corticosteroids [Tables 1 and 2]. Interestingly, the level of transcriptional activation by corticosteroids of elephant shark GR is over 10-fold higher than for elephant shark MR (Figure 3A, B). In this respect, elephant shark GR and MR are similar to human GR and human MR in which cortisol has a higher EC50, as well as a higher level of transcriptional activation for human GR, compared to human MR [15,16,19,20,49].

Activation of elephant shark GR by aldosterone is a notable difference with human GR. Although 10nM aldosterone has little activity for human GR (Figure 3A) [15,27], aldosterone and cortisol have a similar level of activation of elephant shark GR. Aldosterone, the physiological mineralocorticoid in terrestrial vertebrates, also is a weaker activator of truncated skate GR than of truncated skate MR [42] and of lamprey CR [7], which is ancestral to the GR and MR [7,13]. Thus, elephant shark GR retains some responses, albeit diminished, of elephant shark MR and of lamprey CR [7]. We conclude that elephant shark GR is transitional from elephant shark MR towards human GR in specificity for glucocorticoids and mineralocorticoids.

A novel property of elephant shark MR is that it is activated by progesterone and 19norProgesterone [41,50]. Neither steroid activates elephant shark GR. Unexpectedly, progesterone and 19norProgesterone inhibit activation of elephant shark GR by corticosterone (Figure 5A), which suggests that progestins may influence activation of elephant shark GR. Neither progesterone nor 19norProgesterone inhibit activation of human GR by cortisol (Figure 5B). The timing of the loss of recognition of progestins by the GR needs further study.

The evolution of a distinct GR and MR in cartilaginous fish has provided an opportunity to investigate early events in the divergence of these two steroid receptors from their common ancestor. Cartilaginous fish, including elephant shark, also contain a PR and AR, which diverged from a common ancestor. The PR is a close relative of the MR. Comparison of steroid activation of elephant shark PR and AR with each other and with the MR and GR could provide valuable insights into their evolution as important transcription factors in vertebrates.

## Materials and Methods

### Chemical reagents

Aldosterone, cortisol, corticosterone, 11-deoxycorticosterone, progesterone and 19norProgesterone were purchased from Sigma-Aldrich. For the reporter gene assays, all hormones were dissolved in dimethylsulfoxide (DMSO) and the final concentration of DMSO in the culture medium did not exceed 0.1%.

### Construction of plasmid vectors

The full-coding regions and DBD-LBD domains of the GR and MR from *C. milii* were amplified by PCR with KOD DNA polymerase (TOYOBO Biochemicals, Osaka, Japan). The PCR products were gel-purified and ligated into pcDNA3.1 vector (Invitrogen) [51].

### Transactivation Assay and Statistical Methods

Hek293 were used in the reporter gene assay. Transfection and reporter assays were carried out as described previously [51,52]. All transfections were performed at least three times, employing triplicate sample points in each experiment. The values shown are mean ± SEM from three separate experiments. Data from studies of the effect of different corticosteroid concentrations on transcriptional activation of elephant shark GR and MR were used to calculate EC50s for steroid activation of elephant shark GR and MR using GraphPad Prism. Comparisons between two groups were performed using t-test, and all multi-group comparisons were performed using one-way ANOVA followed by Bonferroni test. P < 0.05 was considered statistically significant.

### Genbank Accessions for Domain Comparisons

Elephant shark GR (XP_007899521), human GR (BAH02307), chicken GR (ABB05045), X. laevis GR (NP_001081531), Zebrafish GR (NP_001018547), elephant shark MR (XP_007902220), human MR (NP_000892), zebrafish MR (NP_001093873).

## Author contributions

Y.K., I.M.S., and X.L. performed the research. Y.K., H.U., and S.K. analyzed the data. W.T. and S.H. aided in the collection of animals. Y.K. and M.E.B. conceived and designed the experiments. M.E.B. wrote the paper. All authors gave final approval for publication.

## Data and materials availability

not applicable.

